# Spatial Information in Large-Scale Neural Recordings

**DOI:** 10.1101/002923

**Authors:** Thaddeus R. Cybulski, Joshua I. Glaser, Adam H. Marblestone, Bradley M. Zamft, Edward S. Boyden, George M. Church, Konrad P. Kording

## Abstract

A central issue in neural recording is that of distinguishing the activities of many neurons. Here, we develop a framework, based on Fisher information, to quantify how separable a neuron’s activity is from the activities of nearby neurons. We (1) apply this framework to model information flow and spatial distinguishability for several electrical and optical neural recording methods, (2) provide analytic expressions for information content, and (3) demonstrate potential applications of the approach. This method generalizes to many recording devices that resolve objects in space and thus may be useful in the design of next-generation scalable neural recording systems.

## 1 Introduction

A concerted effort is underway to develop technologies for recording simultaneously from a large fraction of neurons in a brain [1,2]. For a technology to reach the goal of large-scale recording, it must gather sufficient information from each neuron to determine its activity. This suggests that neural recording methodologies should be evaluated and compared on information theoretic grounds, yet no widely applicable framework has been presented that would quantify the amount of information captured by large-scale neural recording architectures.

One common method of identifying neurons in recorded data is through their locations: if the origins of two signals are sufficiently far apart, then they are likely to have come from different neurons. One can formulate this criterion mathematically through the concept of Fisher information, which measures how much information a random variable (e.g. a signal on a detector) contains about a parameter of interest (e.g. the location of origin of that signal). We can only conclude that two neural activity measurements arose from different locations, and thus distinguish the two neurons, if the detected signal contains sufficient Fisher information.

Here we use Fisher information to determine the spatial separability of neural signals. We apply this framework to models of neural recording techniques, describe how the Fisher information scales with respect to recording geometries and other parameters, and demonstrate how this framework can be utilized to optimize experimental design.

## 2 Framework

### 2.1 Localization and Resolution

A foundational concern in neural recording is localization, the ability to accurately estimate the location of origin of neural activity. Localization is a primary method of determining the identity of an active neuron.

The problem of establishing neural locations can be split into two separate regimes. One regime is when an active neuron has no active neighbors (Figure 1A). In this state, we are chiefly concerned with the ability to attribute the signal to the correct neuron (single-source resolution [3]). This can be done by accurately localizing one activity at a given time (Figure 1B&C). The other regime is when two neighboring neurons are simultaneously active (Figure 1D). In this state, we are chiefly concerned with the ability to differentiate the two neurons, i.e. are there two clearly distinguished or one blurred neuron (differential resolution [3]). This can be done by simultaneously localizing the activities of both neurons accurately (Figure 1E&F).^1^

Here we treat both scenarios: first by calculating the Fisher information carried by a single observation about a single neuron's activity, and then by expanding this framework to treat multiple parameters using the matrix form of Fisher information. We address localization in the theoretical limit where the point spread function (PSF) is known, in order to study the limiting effects of neuronal and sensor noise on localization precision.^2^

Regardless of the number of neurons and sensors we are treating, Fisher information gives us a metric with which to evaluate recording. Spatial information, the amount of information regarding the location of a source (i.e., a quantitative measure of localization ability), can be used to determine whether it is possible to correctly attribute an activity to its source (or multiple activities to multiple sources). In order to know the identity of a source, we must be confident about the location of origin of the activity with a positional error less than *δ*, where *δ* is the distance from one neuron to another (Figure 1B&E). In terms of Fisher information, if we have sufficient information to locate the source of activity with a precision *δ*, we can assign that activity to a single neuron that occupies that location.

**Figure 1.**
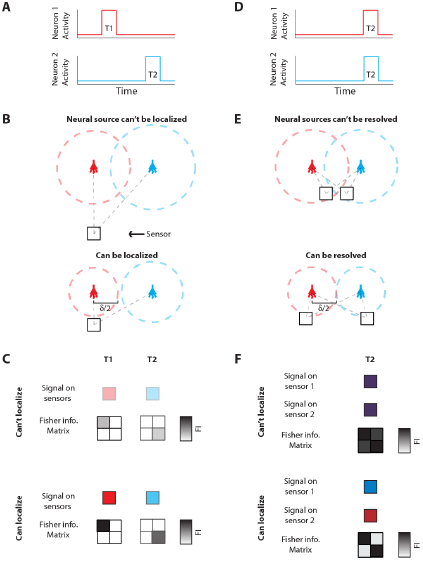
Localization and Resolution. **(A)** In many behavioral states, neural systems have sparse activity, in which neighboring neurons (red and blue) are not active at the same time. In this scenario of single-source resolution, one neuron must be localized at a given time. Panels **B** and **C** look at this scenario. **(B)** Two neighboring neurons are shown a distance *δ* away from each other. Dotted lines indicate regions where we are confident about the source of a signal, i.e. we have a sufficient amount of information regarding that signal’s location. **(C)**The signals from the two neurons are recorded by the sensor at different times and do not interfere with each other. When a neuron cannot be localized effectively, i.e. there is not sufficient Fisher information, it is because the signal from that neuron was not strong enough to overcome noise. **(D)** Sometimes, neighboring neurons are simultaneously active. In this scenario of differential resolution, both neurons must be localized at a given time. Panels E and F look at this scenario. **(E)** Same as **B**, except two sensors are necessary for differential resolution. **(F)** When both sensors record similar signals, i.e. when there is large mutual information regarding the two neurons’ activities, it is difficult to resolve the neurons.

### 2.2 Fisher Information: Single Neuron, Single Sample

We first treat the simplified situation where a single neuron must be localized at a given time, using a single sample (one sensor at one time). This scenario yields simple, intuitive analytic expressions for how the precision of localization scales with a neuron’s location and other experimental parameters. Moreover, activity in neural systems is often sparse [10–14]; thus this simplified scenario may be practical in many cases.

This problem can be approached using Fisher information, *ℐ*(θ), a measure of the information a random variable *X*, with distribution *f(X*; θ) parameterized by θ, contains about the parameter θ [15]:

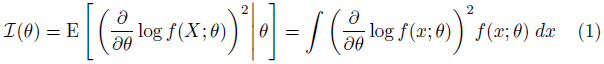

In the context of spatial information in neural recordings, Fisher information quantifies how much the distribution of recorded sensor values *f(X; **d**)* tells us about the location of a signal’s origin with respect to the sensor, **d** (Figure 2D) (here we use boldface to indicate that the location is a vector quantity).

The variance of an unbiased estimator of a parameter is lower bounded by the Cramer-Rao bound (CRB) [16]:

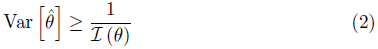

In the context of neural recording, we can use Fisher information to determine how precisely we can estimate the location of activity, given its influence on the sensor output. Note that, while we describe the ability to distinguish neurons solely using spatial information, additional sources of information can be used, e.g., temporal information in optical [17] and electrical recordings [18] (see *Discussion*).

### 2.3 Fisher Information: Multiple Neurons, Multiple Samples

To treat differential resolution (2 active neurons), it is necessary to use multiple samples. In optical techniques, multiple pixels are necessary to localize simultaneously active neurons, and in electrical recordings, multiple electrodes or time-points are required (e.g. a waveform). When we want to know how much information a signal contains about the sources of multiple activities (now ***θ*** is a vector with multiple elements), we must construct a Fisher information matrix with elements:

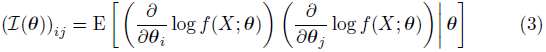

The elements of this matrix represent the information contained about a pair of sources (elements of the vector ***θ***). Note that if sensors have independent noise – an assumption we use for our differential resolution examples – the information matrices can be summed: [*I*(θ)]*_total_* = [*ℐ*(***θ***)]*_sensor_*_1_ + [*ℐ*(***θ***)]*_sensor_*_2_.

The elements of this matrix can be divided into on-diagonal and off-diagonal elements, both of which have practical interpretations. The on-diagonal elements are very similar to the terms for single-unit localization. They describe how much information the ensemble of sensors has about each source. The off-diagonal terms, however, do not have a parallel representation in the single-unit formulation. They can be thought of as crosstalk, or how much the estimation of a source of interest is affected by other sources.^3^

As with Fisher information, the CRB extends to multiple parameter scenarios: the variance of an unbiased estimator of a parameter θ_*i*_ (how precisely a single parameter can be estimated) is:

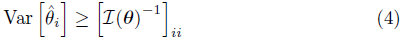

### 2.4 Point Spread Functions and Signal Intensity Distributions

To determine the spatial Fisher information, we must know the distribution of signals on a sensor given the location of the activity, *f*(*X; **d***). The signal measured by many recording systems is well approximated as a linear function of the signals from each neuron in a population [19,20], i.e. the total sensor signal is the sum of the individual neural signals weighted by the magnitude of their individual effects on the sensor (Figure 2A&B). We thus here only consider linear interactions; however, it should be noted that the Fisher information framework is also compatible with nonlinear interactions (e.g. sensor saturation). For *N* neurons and *M* sensors in a system, in the absence of noise, the signal on any particular sensor can therefore be described as:

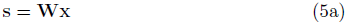

where **s** is the vector of signals on sensors [*s*_1_, …, *s*_M_], and **x** is the vector of signals from neural activities, [*I*_1_, …, *I*_N_]^T^, e.g. the fluorescent signal produced due to neural activity in optical techniques or the voltage signal in electrical techniques. **W** is the matrix of PSFs:

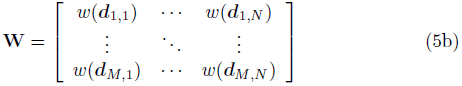

where the PSF, *w*(***d***_*i,j*_), is the scaling of the activity of neuron *j* on the output value of sensor *i*, and is dependent on the location of the activity relative to the sensor (***d***_*i,j*_) and other parameters of a recording modality (e.g. light scattering). The contribution of the activity of neuron *j* to the signal on sensor i is thus *I_j_w*(***d***_*i,j*_).

In the presence of neural and sensor noise, the signal on a given sensor, s_*i*_, is no longer a constant, but characterized by *f_i_(X*), the distribution of signal intensities on a sensor. Here, we assume that the noise can be approximated by a zero-mean Gaussian with variance *σ^2^_noise_,* so that:

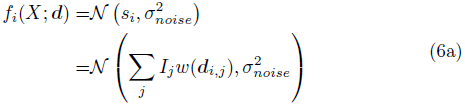

where *𝒩*(μ,σ^2^) signifies a normal distribution (Figure 2C). Focusing on a particular neuron with index *k,* (6a) can be rewritten as:

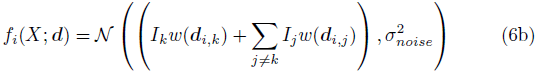

This allows us to calculate the spatial Fisher information in signal *s_i_* for the activity of neuron *k*.

It is important to note that, as long as they can be analytically described, all types of noise (of which there are many; see *Supplementary Information* for further discussion) can be incorporated into this framework. This flexibility in noise sources makes this framework especially relevant for neural recording.

**Figure 2.**
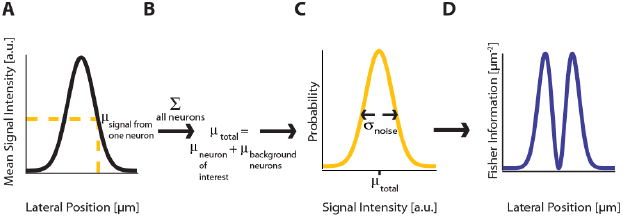
Fisher Information. **(A)** A signal on sensor *i* from a neuron *j* at a particular location has a mean intensity, defined by a recording method’s point spread function and the intensity of the signal from the active neuron. **(B)** The mean total signal on a sensor, *μ_total_*, is the sum of the signals from every neuron. **(C)** The distribution of intensities recorded on a sensor is a function of the total mean signal, *μ_total_*, and the variance of that signal, *σ^2^_noise_*,which can result from many different noise sources. **(D)** Fisher information can be derived from the distribution of signal intensity values on a sensor.

### 3 Applications

Here, we demonstrate the utility of the Fisher information framework for the understanding and advancement of neural recording technologies. We first calculate the spatial Fisher information of both simple and complex recording modalities in the simplified case of a single neuron and single sensor, which reveals insightful scaling properties of these technologies. We next examine a scenario with multiple neurons and multiple sensors in order to determine the extent to which crosstalk between sensors limits localization ability. We finally demonstrate more complex uses of this framework, comparing technologies and optimizing sensor design.

#### 3.1 Analytic Assumptions

We assume that all activity from the neuron of interest, including the noise, is part of the signal of interest. Thus, the total noise is a function of the sensor noise plus the noise of all neurons except for the neuron of interest. The noise will depend on the types of noise and the correlation between noise sources, and we make specific assumptions for our simulations (see below). The assumption that the neuron of interest does not contribute to the noise allows us to provide a simple expression for the spatial Fisher information about activity *k* in a given direction *d^x^* (see *Supplementary Information* for derivation):

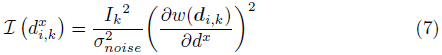

#### 3.2 Simulation Assumptions

For our simulations, we make further assumptions. We emphasize that the following assumptions do not apply to the analytic expressions regarding the scaling of localization ability. In regards to neural activities, we assume that every active neuron has the same activity *I_0_,* while non-active neurons have no activity, that the neuron of interest, *k*, is active at the moment we sample, and that other neurons are active at a uniform rate. We assume noise sources from neurons are independent, so that:

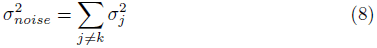

There are many sources of noise, both on neurons and sensors, that could be included, and these are discussed in detail in the *Supplementary Information*. For our applications demonstration, we consider signal dependent noise that can arise from neurons and/or sensors. Specifically, for analytic simplicity, we only consider noise that has a standard deviation proportional to the mean signal: σ^2^_*j*_ ∝ *I*^2^_0_(*w*(***d**_i,j_*))^2^. Under these simplifying assumptions, the magnitudes of the fluorescence (optical) and waveform voltage (electrical) have no influence on the final information theory calculations. We emphasize that these simulation assumptions are implemented to simply demonstrate the use of this framework; more realistic outputs could be found using more complex, realistic noise models.

#### 3.3 Single-source Resolution of Technologies

Here we calculate Fisher information of recording technologies using a single neuron and single sensor to determine technologies’ scaling properties. We look at three technologies: (1) electrical recording, a traditional neural recording modality, (2) wide-field fluorescence microscopy, a traditional optical approach, and (3) two-photon microscopy, a modern optical approach. These examples are chosen for their relative simplicity and ability to illustrate the flexibility of a Fisher information approach to modeling neural recording.

We calculate Fisher information in the most meaningful direction for the techniques we consider: radial for electrical recording (due to spherical symmetry), and lateral for the optical techniques (due to cylindrical symmetry). For electrical recording, the origin is at the center of measurement (at the electrode). For optical recording, the origin is at the center of the lens. For any technology, the aim is for there to be, across all sensors, sufficient information about every location in the brain in order to identify a neuron firing in that location.

Thus for an individual sensor, it can be better to have sufficient (enough to identify a neuron, as in Figure 1) information spread over a large area than excessive information about a small area. This suggests that experimental designs could be modified to get sufficient information for the required task. For example, an optical technology may have extra information at low depths, but insufficient information at large depths. In this case, the PSF could be modulated (e.g. [21]) to decrease low-depth information (making those images blurrier), while increasing high-depth information.

#### 3.4 Electrical Sensing

The electrical potential from an isolated firing neuron decays approximately exponentially with increasing distance [22,23], at least at short distances. Here, we model a simple electrical system: an isotropic electrode with spherical symmetry. In this isotropic approximation, the PSF has an exponential decay with radial distance from the electrode tip (Table 1; Figure 3B).

For electrical recording, spatial Fisher information (derived from the above PSF) decreases exponentially with increasing distance from the electrode (Table 1, Figure 3C&D). Given this behavior and the properties of electrical signals in the brain, electrical recordings provide relatively weak information over a relatively wide area.

**Figure 3.**
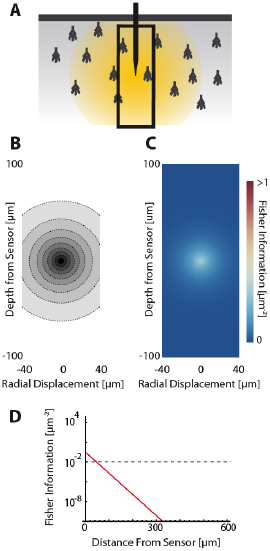
Electrical Recording. An overview of the modeling and Fisher information analysis of electrical recording. **(A)** Schematic: An electrode records electrical signals directly from nearby neurons. The black box indicates the space represented in panels **B** and **C**. **(B)** The spatial PSF for electrical recording, valued in arbitrary units, for an electrode located at (0,0). **(C)** The radial Fisher information contained about signals from different locations within a volume surrounding the electrode in panel B. **(D)** The Fisher information for a point with a given distance from the electrode in panel **B**. The grey dashed line indicates the minimum Fisher information needed for a CRB standard deviation of 10 μm. This 10 μm standard deviation corresponds to a 95% accuracy of determining the correct active neuron for neurons whose centers are 40 μm apart, and assuming a Gaussian estimation profile.

#### 3.5 Optical Sensing

##### 3.5.1 General Information

Optical recording of neural activity generally relies on fluorescent dyes that are sensitive to activity. In order to measure this signal, a neuron must be illuminated with light in the dye’s excitation spectrum. Light is then emitted by the dye at a distinct, longer (lower energy) wavelength, which is picked up by a photodetector. Optical signal transmission is subject to absorption, scattering, and diffraction, which degrade the emitted signals with distance. Absorption of light effectively cause an exponential decrease in intensity of detected photons as light travels through a medium [24,25]. Scattering can affect light in multiple ways; high-angle scattering diverts photons from the detector and produces an effect similar to absorption, while low-angle scattering causes blurring of the image on the detector. This blurring increases approximately linearly with depth into the tissue [26]. Finally, diffraction results when light passes through an aperture, creating the finite-width Airy disk [27]. In our optical PSFs, we assume scattering and diffraction result in Gaussian blurring [26,28]. Our PSFs assume imaging through a single homogeneous medium; in practice, tissue inhomogeneity and refractive index mismatch can produce additional aberrations in the absorption, scattering, and diffraction domains that we do not model here.

In a typical optical setup, a lens focuses a set of photons from one point in space onto a corresponding point behind the lens. This phenomenon can be used either to focus incident light onto a desired location for illumination, or to focus emitted light from the focal plane onto a photodetector for imaging. Photons from outside the focal plane will be blurred, and this blurring increases linearly as distance from a focus point increases [29,30]. We also assume defocusing results in Gaussian blurring [29,30].

##### 3.5.2 Wide-field Fluorescence Microscopy

Neural activity in a focused optical system is generally sensed using fluorescent dyes, which require some excitatory light. In the canonical optical example of wide-field microscopy, an entire volume is illuminated (Figure 4A). The PSF for this technology takes the above effects of absorption, scattering, diffraction, and defocusing into account; we assume total illumination so that the PSF here models the spread of the emission light (Figure 4B, Table 1).

For optical recording with a simple lens, Fisher information is concentrated on a ring around the axis of the lens within each focal plane (Figure 4C). This stems from the fact that slight deviations from the imaging axis do not change the signal intensity, and thus there is no location information contained in the signal intensity at those locations. For large depth, the ability to distinguish locations decreases exponentially due to photon loss caused by scattering and absorption (Table 1). For medium depth ranges, scattering blurs the image, even on the focal plane. The decrease in localization ability from this effect follows a fourth order polynomial.^4^ Defocusing produces a similar information loss outside of the focal plane. Thus optical recordings provide a large amount of information on the focal plane and this information, largely due to scattering, rapidly decreases as the distance from the lens increases (Figure 4D).

**Figure 4.**
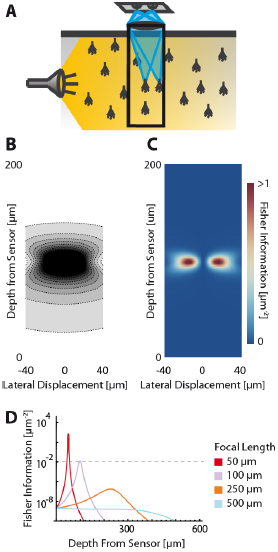
Wide-field Fluorescence Optical Recording. An overview of the modeling and Fisher information analysis of wide-field fluorescence optical recording. **(A)** Schematic: The whole recording volume is illuminated; dye in active neurons fluoresces and emits light; the emitted light is focused by a lens onto a photosensor. Here, we analyze one point on that photosensor. The black box indicates the space represented in **B** and **C**, with zero depth being located at the lens. **(B)** The spatial PSF for wide-field fluorescence optical recording, valued in arbitrary units, for a lens centered at (0,0) with a focal plane at 100μm. **(C)** The lateral Fisher information contained about signals from different locations within a volume sensed by the wide-field fluorescence optical system shown in **B**. **(D)** The maximum Fisher information for a point with a given depth from the sensor for systems with focal planes at different depths. The grey dashed line indicates the minimum Fisher information needed for a CRB standard deviation of 10 μm.

##### 3.5.3 Two-photon Microscopy

In two-photon microscopy, long-wavelength incident light (i.e. composed of low-energy photons) is focused onto a single point of interest to excite fluorophores in that area. In order for the fluorophore to emit light, two low-energy photons must be absorbed nearly simultaneously; the likelihood of this event is proportional to the square of the intensity of incident light at a point. Effectively, this concentrates the area of sufficient illumination to a volume nearby the focal point of the incident beam (while increasing the illumination power requirements) [31]. Like with wide-field fluorescence microscopy, the PSF is a function of defocusing, absorption, and scattering (Figure 5B, Table 1). We assume total photon capture so that the PSF here models the spread of the excitation light.

For two-photon microscopy, Fisher information is also concentrated in a ring around the beam axis, and it is more tightly concentrated than in wide-field fluorescence microscopy (Figure 5C). This is due to the reduced scattering of longer wavelengths of incident light, as well as decreased fluorescent excitation of laterally-displaced neurons. The ability to estimate a neuron’s location still scales with the standard deviation to the 4^th^ power (Table 1).

**Figure 5.**
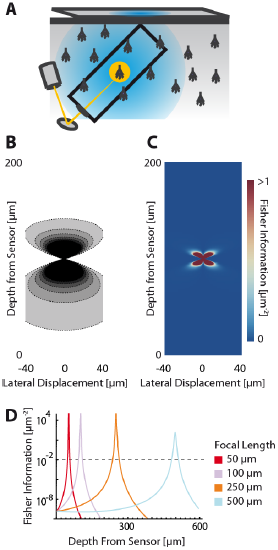
Two-photon Optical Recording. An overview of the modeling and Fisher information analysis of 2-photon optical recording. **(A)** Schematic: incident light is focused onto a particular location in a volume; dye in neurons illuminated by the incident light fluoresces and emits light; the emitted light is sensed by a large single photosensor. The black box indicates the space represented in **B** and **C**, with zero depth being located at the lens and increasing depth indicating increasing distance into the brain. **(B)** The spatial PSF for incident light relative to its source in 2-photon optical recording. It is valued in arbitrary units for a lens centered at (0,0) with a focal plane at 100 μm. **(C)** The lateral Fisher information contained about signals from different locations within a volume sensed by the 2-photon optical system shown in panel **B**. **(D)** The maximum Fisher information for a point with a given depth from the light source for systems with focal planes at different depths. The grey dashed line indicates the minimum Fisher information needed for a CRB standard deviation of 10 μm.

**Table 1.**
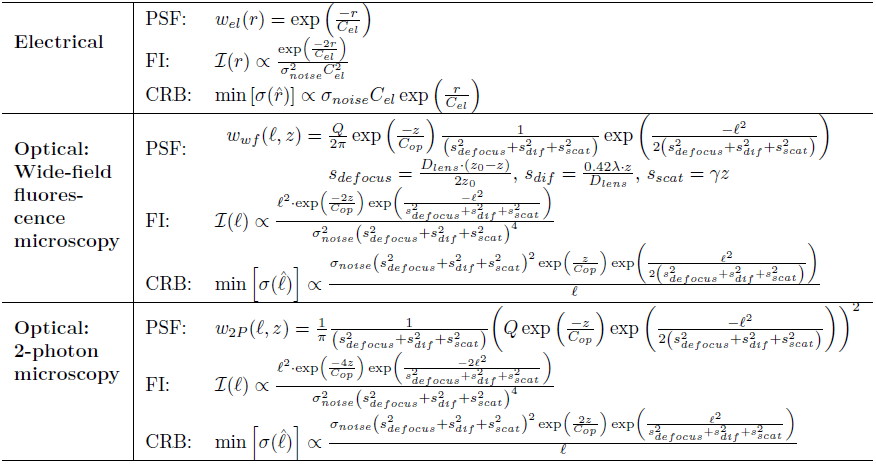
Point Spread Function, Fisher Information, and Cramer-Rao Bounds of Recording Modalities. Analytic expressions are given for PSFs, and for Fisher information and CRB proportionality. Fisher information is in the most meaningful direction for the given technology: *r* is the distance in any radial direction, and *𝓵* is the lateral distance for optical techniques. *C_el_* is the spatial constant of electrical decay. *C_op_* is the spatial constant of optical decay. *s*^2^*_defocus_*, *s*^2^_*dif*_, and *s^2^_scat_* are the variance of the spread of optical light due to de focusing, diffraction, and scattering, respectively. *D_lens_* is the diameter of a lens. λ is the wavelength of the light. *z*_0_ is the focus depth, and *Q* is the light flux (area per photon).

**Table 2.**
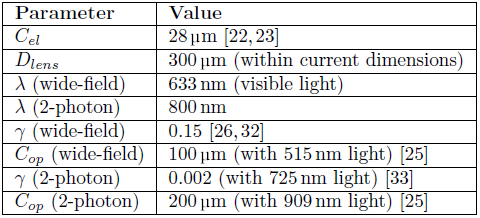
Simulation Parameter Values

| Parameter | Value |
| --- | --- |
| $C_{el}$ | 28 $\mu\text{m}$ [22, 23] |
| $D_{lens}$ | 300 $\mu\text{m}$ (within current dimensions) |
| $\lambda$ (wide-field) | 633 nm (visible light) |
| $\lambda$ (2-photon) | 800 nm |
| $\gamma$ (wide-field) | 0.15 [26, 32] |
| $C_{op}$ (wide-field) | 100 $\mu\text{m}$ (with 515 nm light) [25] |
| $\gamma$ (2-photon) | 0.002 (with 725 nm light) [33] |
| $C_{op}$ (2-photon) | 200 $\mu\text{m}$ (with 909 nm light) [25] |

#### 3.6 Differential Resolution versus Single-source Resolution

We now extend the Fisher information framework to the case of multiple sources and multiple sensors. Previously, when considering a single source, localization ability was only limited by noise. With multiple sources, resolution ability is additionally limited by crosstalk between sensors and confounding sources. We ask how the ability to localize neurons differs between the previously discussed sparse firing scenario, and the scenario in which multiple neurons are simultaneously active. That is, how will the previously derived expressions be hindered due to crosstalk? In addition, we look at the benefits imparted on localization by additional sensors.

Here we consider a case where there are two sensors, each of which is located below a neuron (Figure 1E).^5^ This setup is analogous to most optical setups, where a single pixel corresponds to a small area that the pixel is supposed to image. We first look at Fisher information contained about each source by the two-sensor ensemble, when there is no crosstalk (noise is the only limit). This is equivalent to looking at the Fisher information that the two sensors contain about a single source. As expected, as sources move further away from sensors, the CRB for the two-sensor case scales similarly to the previously seen single-sensor case (Figure 6A). The primary difference is that we are using two sensors instead of one and assuming additive information. Thus, there is slightly more information from the two-sensor setup than the one-sensor setup, resulting in slightly tighter CRBs for the two-sensor ensemble.^6^

Finally, we specifically look at the effect of crosstalk on the ability to estimate neurons’ locations (Figure 6C). We define crosstalk as the ratio of the signal received from the confounding source to the signal received from the primary source (i.e. the ratio of off-diagonal to diagonal elements in the Fisher information matrix). The system is relatively robust to small amounts of crosstalk: the CRB increases only 10% for a crosstalk of 0.3. However, there are profound effects when there are large amounts of crosstalk: for a crosstalk of 0.9, the CRB increases by over 400%. It is important to note that excellent resolution is possible with high crosstalk, and poor resolution is possible with low crosstalk. For instance, while there may be high crosstalk at the focal plane, the large increase in the CRB still yields acceptable resolution (Figure 6D). Conversely, while there may be low crosstalk away from the focal plane, there is nonetheless poor resolution.

**Figure 6.**
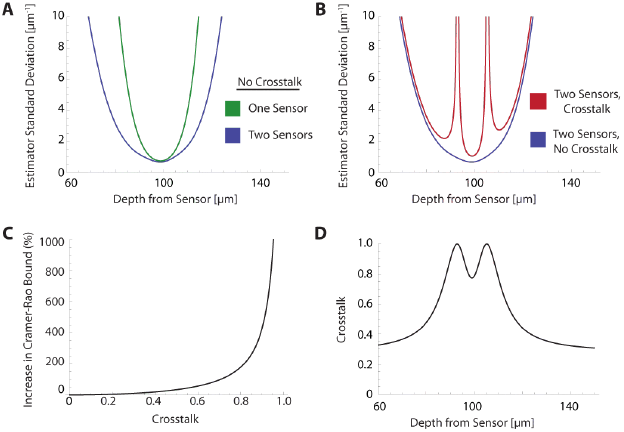
Localization with Multiple Sources. For all panels, we simulate two sensors 40 μm apart and two sources 30 μm apart centered between the sensors using a wide-field fluorescence microscopy setup with a focal length of 100 μm. Both sources are located in the same plane. We look at how CRBs on the estimation of both sources vary as the two sources change depth with respect to the sensors (note that the CRBs for each source are equal due to the symmetric setup). **(A)** A comparison of CRBs on a location estimate without crosstalk using one sensor vs. two sensors assuming additive Fisher information. **(B)** A comparison of CRBs on a location estimate by two sensors when signals are allowed to interfere with each other (crosstalk) vs. a hypothetical situation with no interference (no crosstalk). **(C)** The effect of crosstalk (here defined as the ratio of the signal received from the confounding source to the signal received from the primary source), on the increase in the CRB regarding estimation of neuronal locations. **(D)** The amount of crosstalk in panel **B** as a function of the neurons’ depths from the sensor.

#### 3.7 Technology Comparisons

In order to determine appropriate technologies for a given situation, it is necessary to know which technology will maximize the information output, and where information will be concentrated for a given technology. Here we apply this Fisher information framework to a two-source, two-sensor setup for both wide-field fluorescence and two-photon microscopy in order to determine performance over depth (Figure 7). We find, perhaps confirming intuition, that wide-field and two-photon fluorescence perform similarly for shallow sections, but performance of wide-field fluorescence microscopy degrades significantly at a depth of 500 μm while two-photon performs well at this depth. Interestingly, both methods contain a large amount of information not only about signals near the focal point, but also about sources nearby the lens. This implies that signals could be recovered from out-of-focus samples given proper recording conditions. While this demonstration yielded the expected results, this framework could be used to compare existing technologies in novel situations, or to compare novel technologies.

**Figure 7.**
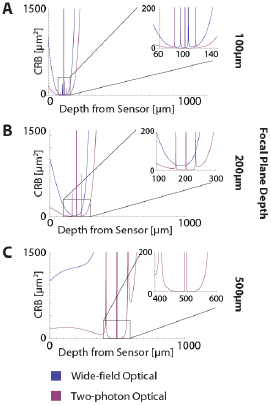
Optical Technology Comparison at Multiple Focal Depths. CRB on the location of a single source in a two-sensor, two-source system with crosstalk. The sensors are 40 μm apart and the sources are 30 μm apart situated between the sensors. The depth of the sources is varied by an equal amount and the CRB on each of the sources is calculated at each depth (the CRB of each source is equivalent due to the symmetric setup). This analysis is performed for wide-field fluorescence and two-photon optical systems with focal depths of **(A)** 100 μm, **(B)** 200 μm, and **(C)** 500 μm. Insets provide detail about behavior of optical systems at locations near the desired focal depth.

#### 3.8 Sensor Placement

In order to successfully record activity from every neuron in a volume, we must place sensors so that they extract sufficient information about every neural location in that volume. That is, the tiling of the Fisher information from every sensor needs to cover the entire volume. Mathematically, the Fisher information contained about each point in a volume must exceed some threshold for localization. With this framework, it is possible to ask questions about the necessary experimental parameters of neural recording technologies.

Here, we simulate several possible arrangements of electrical sensors and evaluate the Fisher information that these systems provide about different locations in a volume. Specifically, we look at four electrode arrangements: (1) columns of electrodes where electrodes are densely packed within a column, and these columns are arranged in a grid [34] (Column electrodes); (2) randomly placed electrodes (Random electrodes); (3) electrodes evenly distributed in a plane (Planar electrodes); and (4) electrodes evenly distributed in an equilateral grid (Grid electrodes) (Figure 8A). Here, we assume that noise is independent between sensors, i.e. noise is all on the sensor. Under this assumption, each electrode takes an independent sample of a signal; information about the location of the source of that signal is then additive across sensors. Fisher information here is thus the information the entire ensemble of electrodes provides about a point. For simplicity of this demonstration, we use the single-source resolution formulation of these scenarios, which corresponds to the common situation of sparse neural firing. Crosstalk between sources on sensors would necessarily reduce the amount of information contained about individual sources and would be geometry-dependent. The distribution of Fisher information within a volume varies based on electrode placement (Figure 8B), with Fisher information being highly concentrated in areas with high electrode density.

Just as we want to know where systems concentrate Fisher information, we want to know how often we will have sufficient spatial Fisher information. To do this, we look at the distribution of Fisher information within a volume (Figure 8C). In this simplified simulation, columnar electrodes have a distribution that sufficiently estimates most locations: a small fraction of points lie in between electrodes and their location cannot be estimated well, while the significant fraction that lies close to a column can. Random electrodes have sufficient information about all locations in the simulated volume. Planar electrodes have a log-constant distribution; they carry very little information about most locations in a volume, but carry a large amount of information about a small fraction of locations. The use of Grid electrodes has comparable performance to Random electrodes in that it carries sufficient information about all locations in a volume. Due to the regular nature of Grid electrodes, there is the added benefit of a guaranteed lower bound for information carried about locations in a volume. The use of this Fisher information framework promises to inform sensor placement decisions.

**Figure 8.**
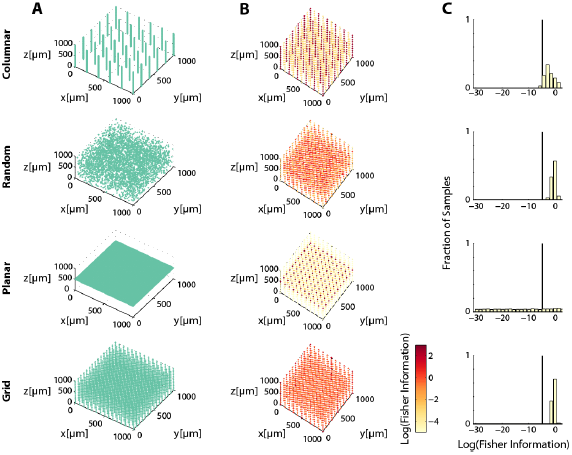
Electrode Placement and Fisher Information. Simulation of ~ 3.5 × 10^3^ electrodes in a 1 mm × 1 mm × 1 mm cube of brain tissue. Electrodes were arranged in one of four patterns: a 6 × 6 grid of columns of electrodes with 100 electrodes evenly distributed in each column (top row), random placement (second row), electrodes uniformly distributed on a plane at 500 μm depth (third row), and uniformly distributed in a grid throughout the volume (bottom row). Total Fisher information about a point consists of the sum of information contained about that point in each sensor. This analysis considers information about radial location. **(A)** Distribution of electrodes in the volume for each pattern. **(B)** Total Fisher information contained about 10^3^ uniformly distributed points in the volume. Each point is displayed at its location in the volume, and its color indicates the amount of Fisher information. Fisher information is plotted with a floor of 10 × 10^−5^ μm^−2^. **(C)** Distribution of Fisher information contained about a random sample of 10^4^ points from the volume. Vertical black lines indicate Fisher information corresponding to a 10 μm CRB on standard deviation. See Table 2 for parameter values.

### 4 Discussion

Here we have developed a framework to quantitatively think about the challenges of large-scale neural recording and determine the necessary experimental parameters for potential recording modalities. This extends previous work applying Fisher information to individual imaging techniques (e.g. [9,21,35-41]) by considering a PSF that models recording in neural tissue, and then using the CRB to establish signal separability rather than spatial resolution. We have demonstrated the utility of this framework by characterizing the scaling properties of common recording techniques, and then showing how experimental design can be influenced by information considerations.

We made a large number of simplifications when demonstrating the use of the Fisher information framework. Our modeling of (1) neural activities and noise and (2) the details of the recording methods is significantly simplified. (3) Many recording methods allow for ways of multiplexing – of using each sensor to get multiple pieces of information – which we do not consider. (4) This framework is solely focused on distinguishing neurons using spatial information, while other information sources may be used. However, these approximations were useful in demonstrating a unifying view over recording methodologies in a single paper. Applying this framework with a smaller set of simplifying assumptions would offer more precise insights. We will discuss such potential developments for each of the four aforementioned simplifications.

First, we asked how we could use recording channels to identify the location of a fixed, known, activity. In practice these activities fluctuate over time, and can differ based on spike type (e.g. simple vs. complex in Purkinje cells). Interestingly, it would be possible to use Fisher information to differentiate spike types given a single source recording. Moreover, the various noise sources were approximated by a simple function that ignores many potential sources of noise (see *Supplementary Information*). A comprehensive model of noise affecting neurons and sensors does not yet exist. Further research in this area will yield more informative results.

Second, we asked how we could use simplified models of recording systems to estimate the locations of neurons. However, the models of the recording channels are likely overly simplified. For example, for optical recordings we assumed scattering through homogenous tissue, and for electrical recordings we ignored the filtering properties of electrodes. There exists a rich literature of modeling optical and electrical systems that could allow better models of recording modalities (e.g. [25,42]); incorporating these models into the framework may alleviate some of the concerns over oversimplification, and may even provide a framework for validating those models.

Third, while we have assumed that sensors only have access to an instantaneous signal amplitude, almost all recording methods have sensors that provide different information over time, a kind of multiplexing. Thus, additional information can be gained by combining information across time. Multiplexing, both from temporally distinct measurements and other methods, can be readily included in the Fisher information framework by introducing virtual recording channels.

Fourth, signals from different neurons need not solely be separated using spatial information. In fact, this may be necessary for certain recording situations, e.g. where the dendrites of one neuron produce a signal within the CRB of the cell body of another neuron. In order to determine the neural, rather than spatial, sources of these signals, extra information would be needed. There are several methods to encode additional information. For example, if each neuron could produce optical activity at a different, identifiable, frequency, then it would be possible to record from all neurons without knowing the locations of the neurons (similar to [43]). Similar considerations exist for electrical waveforms of spikes and many other techniques. For example, molecular recording devices, which store neural signals in macromolecules such as DNA [44], could identify neurons through the barcoding of molecular DNA [45]. Nonetheless, spatial information is one of the common ways of separating neural activities within a channel and is effective at doing so.

The framework that we have introduced here is not limited in application to the recording methods described. For instance, there are many modern optical techniques being used for large-scale recording [46,47]. Additionally, both ultrasound and MRI have been proposed as potential recording media for large-scale neural recording [2,48]. With a proper PSF describing how signals from different positions in the brain reach a sensor (some discussion in [47,49–53]), this framework could easily be applied to determine bounds on signal separability for those techniques. Likewise, this framework could be used to analyze methods that have not yet been described, and thus promises to be a useful tool in the advancement of neural recording technologies.

Fisher information, in the context of PSFs that define how neurons affect recorded channels, allows a unifying view of currently existing recording methods. It promises a way to optimize existing technologies and estimate potentials of proposed technologies. We believe that a Fisher information framework can be a useful tool for the emerging field of scalable recording techniques.

## 5 Additional Methods

### 5.1 Noise Calculations

In our Applications simulations, we make several assumptions about noise. We assume noise sources are uncorrelated (i.e. the noise from each neuron is independent and independently distributed). The sensor signal variance arises from signal dependent noise, with a standard deviation proportional to the mean signal. The signal dependent noise can be on all background neurons and/or on the sensor. As the mean activity is *I*_0_, the standard deviation of the activity is *α · I_0_*, where α is a constant. The activity that reaches the sensor *i* (the signal) from a given neuron *j* then has a variance of σ^2^_j_ = α·(Ι_0_ · *w(**d**_i,j_*))^2^. As the noise sources are independent, their variances can be added, so 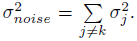 We assume that neurons are uniformly distributed across the brain with density *ρ*_*space*_ and that all neurons have the same probability of firing at a given time, *ρ*_*fire*_.

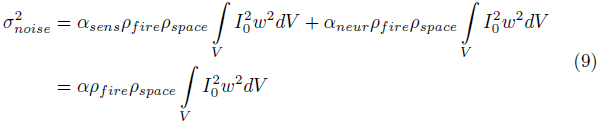

In our simulations, we set *α* = 0.1 (action potentials have SNRs ranging from 5-25 [54]), *ρ*_*fire*_ = 0.01 (assuming neurons on average fire at 5Hz [55] and action potentials last ≈ 2ms), and *ρ*_*space*_ = 67000 mm^−3^ (dividing the number of neurons in the human brain, ≈ 8 × 10^10^ [56] by its volume, ≈ 1200 cm^3^ [57]).

### 5.2 Applications: Electrode Grid Analysis

Electrode locations were assigned to nodes on a 1μm grid spanning a 1 mm × 1 mm × 1 mm cube using the following procedures:

*Columnar*: Column locations were spaced evenly, 200 μm apart, on a 6 × 6 grid in the x-y plane. 101 electrodes were distributed evenly along each column, 10 μm apart.

*Random:* Locations on the grid were drawn from a uniform random distribution with replacement.

*Planar*: Electrodes were placed on a uniform 61 × 61 grid in the x-y plane, corresponding to a grid spacing of 17 μm, with a depth of 500 μm.

*Grid*: Electrodes were placed on a uniform 15 × 15 × 15 grid in the volume, corresponding to a grid spacing of 71 μm.

These procedures give locations for 3636, 3636, 3721, and 3375 electrodes respectively.

For each arrangement of electrodes, we determine the total Fisher information that the ensemble of points contains about some point, assuming independence of noise between sensors:

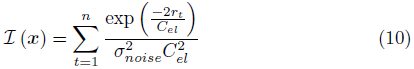

Where *x* is the point we calculate the Fisher information about, *n* is the number of electrodes, and *r_t_* is the Euclidian distance between *x* and the *t*-th electrode.

## 6 Acknowledgements

We would like to thank Dario Amodei and Darcy Peterka for their helpful comments. We would like to thank Dan Dombeck for helpful discussions regarding optics and Mikhail Shapiro for discussions regarding MR applications.

Adam Marblestone is supported by the Fannie and John Hertz Foundation fellowship. Konrad Kording is funded in part by the Chicago Biomedical Consortium with support from the Searle Funds at The Chicago Community Trust. Konrad Kording is also supported by NIH grants 5R01NS063399, P01NS044393, and 1R01NS074044. Bradley Zamft is supported by the American Association for the Advancement of Science Science and Technology Policy Fellowship. George Church acknowledges support from the Office of Naval Research and the NIH Centers of Excellence in Genomic Science. Edward Boyden acknowledges funding by Allen Institute for Brain Science; AT&T; Google; IET A. F. Harvey Prize; MIT McGovern Institute and McGovern Institute Neurotechnology (MINT) Program; MIT Media Lab and Media Lab Consortia; New York Stem Cell Foundation-Robertson Investigator Award; NIH Director’s Pioneer Award 1DP1NS087724, NIH Transformative Awards 1R01MH103910 and 1R01GM104948, NSF INSPIRE Award CBET 1344219, Paul Allen Distinguished Investigator in Neuroscience Award; Skolkovo Institute of Science and Technology; Synthetic Intelligence Project (& its generous donors).

## 8 Supplementary Information

### 8.1 Noise Sources

The Fisher information framework allows for arbitrary noise sources, so long as they are able to be modeled. However, to demonstrate potential applications, we used a very simplified noise model that only considered signal dependent noise where the standard deviation was proportional to the mean.

There are multiple potentially relevant sources of noise that could readily be included in our model. (1) Each sensor has a constant level of noise simply due to thermal effects. (2) Many sensors have an additional variance that is proportional to the signal, e.g. due to low numbers of photons (shot noise). (3) Many sensors have an additional variance that is proportional to the square of the signals, e.g. reference fluctuations. (4) Each neuron may produce constant noise, e.g. background fluorescence of dyes. (5) Each neuron may produce variance that linearly depends on its signal strength, e.g. fluorophore activations. (6) Each neuron may produce variance that quadratically depends on its activation, e.g. action potentials that propagate back into varying parts of the dendritic tree.^7^ Plus, these noise sources may be independent across neurons or correlated. We have some knowledge about the exact sizes of these signals [2], but most of these numbers are hard to know. They may be reasonable to measure in future experiments.

Taking these signals together, we obtain the following noise level on a sensor *i* (given a recording of *N* firing neurons indexed by *j*):

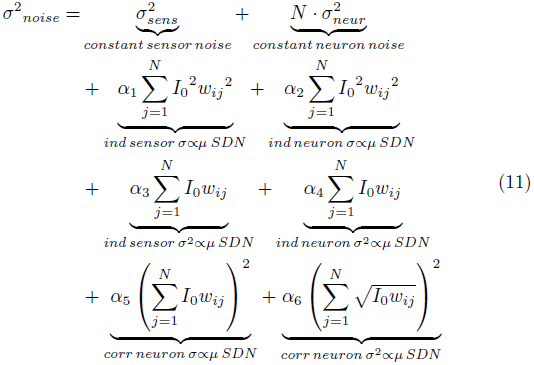

where *ind* and *corr* refer to independent and correlated noise sources, and *SDN* refers to signal dependent noise.

Assuming, as we do in the main text, that neurons are uniformly distributed and have a uniform firing rate across the entire volume:

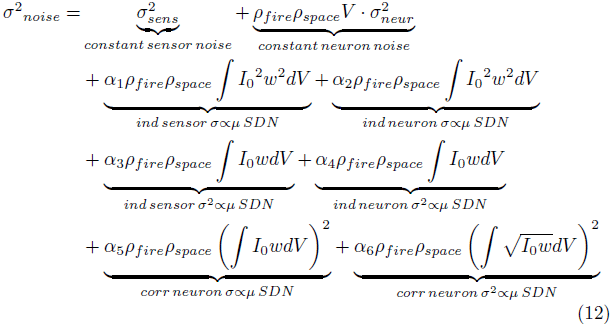

Both constant and shot noise terms can be minimized in their effect by optimizing the experimental design, e.g. through good dyes and strong illumination (but see [2]).

In addition, in the main text we assume that the noise is Gaussian, which has also been assumed in previous statistical formulations [9,38]. This assumption has been shown to be valid for thermal noise and shot noise in some conditions [58,59].

### 8.2 Fisher Information Derivation

We have a distribution 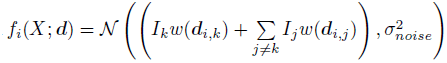 (See (6b)) and are interested in finding its Fisher information with respect to a given direction *d^x^.* For simplicity of notation, we let 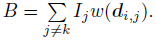

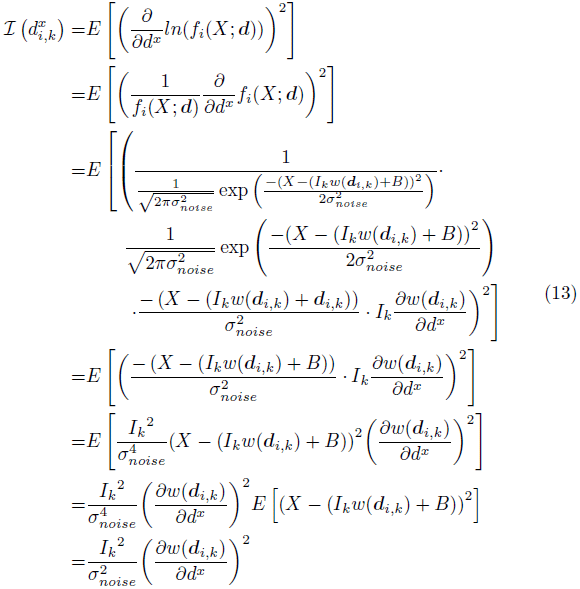

This results in (7). The above derivation will hold for any distribution with zero-mean Gaussian noise.

## Footnotes

1 While we have been discussing differentiating neurons, the framework itself differen-tiates between point sources. In this paper, we make the assumption that separate point sources belong to separate neurons. In reality, it is possible that there could be separate signals from the cell body and dendrites that are perceived as different sources. These can be united using additional information (e.g. anatomical imaging or simultaneous activity).

2 There exists a family of deconvolution techniques that estimate the PSF and use it to obtain a more accurate representation of the original signal (e.g. [4–7]). In theory, with sufficient samples and knowledge of the PSF, one could obtain a perfect representation of a sparse signal in the absence of noise. This is not the case in practice, as signals are not only modified reversibly by PSFs, but are modified irreversibly by noise on neurons and detectors (e.g. [8,9]). In the presence of noise and other aberrations, it thus becomes difficult to isolate individual sources using deconvolution techniques, even when the PSF is known. Thus, it is interesting to determine the isolated effects of noise on recording methods.

3 From a physics perspective, the off-diagonal term (*ℐ*(θ))*ij* describes the coherence between the signals from sources *i* and *j*. The diagonals are the incoherent terms.

4 A curious implication here is that the standard deviation of position estimation is proportional to the fourth power of the standard deviation of possible locations, rather than being simply linearly proportional as one might expect. This is a result of using signal intensity as a proxy for location: information about location is proportionate to the square of how much signal intensity changes as location changes (i.e. ℐ ∝ (ω′)^2^). It is also a function of our Gaussian noise assumption. Given a 2-D distribution of possible locations for a signal’s source, the change in illumination between different points in that distribution scales with the spread of that distribution to the fourth.

5 Sources in this formulation are laterally offset slightly from a corresponding sensor. Here, the offset is 5 μm. This is done in response to sensors containing no Fisher information about a source directly above it, as has been discussed above.

6 It is important to note that these are not discontinuities, as crosstalk never equals or exceeds the desired signals (e.g. see Fig. 7, which shows a greater range).

7 In a simplistic model, when a neuron fires, the action potential spreads into some variable proportion of the dendritic tree. If the recorded signal is dependent on the proportion of dendritic branches the action potential propagates into, then the standard deviation of the recorded signal is proportionate to the mean signal entering the dendrites.

